# Cytokinesis genes regulate ring canal patterning and follicle cell differentiation in *Drosophila*

**DOI:** 10.64898/2026.09.10.750734

**Authors:** Cristy P. Mendoza, Andreana Gomez, Michael Baumgartner, Todd G. Nystul

## Abstract

Stable intercellular bridges that form through incomplete cytokinesis are present in a wide variety of cell types but their function in somatic cell differentiation is not well understood. Using super-resolution confocal microscopy, we identified a stepwise process of ring canal development in the follicle cells of the *Drosophila* ovary. Using a custom-trained deep learning model to aid in 3D image segmentation and follicle cell quantification, we found that ring canals are more heterogeneous at the early prefollicle stages compared to the more differentiated later stages. In addition, we found that depletion of the septin, *peanut*, caused cytokinetic defects but did not disrupt ring canal formation, whereas depletion of the ESCRT III gene, *shrub*, increased ring canal number in prefollicle cells and caused proliferation and differentiation phenotypes. Our findings identify cytokinesis and ring canal formation as a point of regulation in the patterning of proliferation and differentiation in the *Drosophila* follicle stem cell lineage.

**Summary:** Somatic ring canals in the *Drosophila* ovary develop through a stepwise series of states and are a point of regulation of proliferation and differentiation toward a postmitotic cell fate called stalk cells.

## Introduction

In stem cell-based tissues, cell fate specification must be precisely patterned to ensure that stem cells and their daughters acquire the appropriate identities at the correct time and place to support tissue function. This complex orchestration of events is achieved through the integration of inputs from multiple sources that regulate the necessary changes in gene expression, including both the signal transduction cascades that are the direct regulators of transcription factors and many aspects of cell biology, such as actin dynamics, mechanical forces, intracellular pH, and metabolism (Chacón-Martínez et al., 2018; Tatapudy et al., 2017; De Belly et al., 2022). A wide range of studies have demonstrated that these aspects of cell biology can serve as both repositories for the storage of information about cell states and regulators that drive changes in cell behavior.

One well-studied aspect of cell biology that is emerging as an important contributor to the regulation of proliferation and differentiation in stem cell lineages is cytokinesis (Mathieu et al., 2022; Thieleke-Matos et al., 2017). During cytokinesis, an actomyosin contractile ring drives the initial stages of cleavage furrow ingression and then transitions into a midbody ring that typically resolves to complete abscission, resulting in two separate sister cells (Steigemann and Gerlich, 2009). However, the timing of abscission can vary, often occurring several hours after mitotic exit. In some cases, the midbody ring is stabilized, leaving an intercellular bridge between the sister cells that allows for the continued exchange of cytoplasmic material between sister cells (Fawcett et al., 1959; Burgos and Fawcett, 1955; Koch and King, 1966). These intercellular bridges have been found in a diverse range of organisms, from invertebrates such as insects and hydra to mammals (Chaigne and Brunet, 2022; Price et al., 2023). They are particularly common in germ cell lineages, where they contribute to the coordination of mitosis, meiosis, and other critical processes by facilitating the intercellular transport of nutrients, macromolecules, and organelles (Robinson et al., 1994; Haglund et al., 2011; Cox and Spradling, 2003; Sorkin et al., 2025). In addition, intercellular bridges or delayed cytokinesis has also been observed in some somatic cell types, including cells in the early mammalian embryo, HeLa cells, and ovarian granulosa cells, though much less is known about how intercellular bridges form and function in somatic cells (Doherty et al., 2022; Komatsu and Masubuchi, 2018; Zenker et al., 2018; Estey et al., 2010).

The *Drosophila* ovary has been a highly informative model for the study of both germline and somatic intercellular bridges, which are referred to as “ring canals” (Price and Lewellyn, 2026; Airoldi et al., 2011; McLean and Cooley, 2013). Each ovary is composed of long strands of developing follicles, called an ovariole, and new follicles arise from a structure at the anterior tip of each ovariole called the germarium (**Fig. 1A**) (Spradling et al., 1997; Kirilly and Xie, 2007). Within the germarium, 2-3 germline stem cells divide asymmetrically to self-renew and produce daughter cells called cystoblasts. Cystoblasts then undergo four rounds of mitosis with incomplete cytokinesis, forming a 16-cell cyst in which neighboring cells are connected by stable ring canals. When these cysts move into the posterior half of the germarium, they become surrounded by a layer of epithelial follicle cells produced by follicle stem cells (FSCs). FSC daughter cells differentiate into prefollicle cells (pFCs) that then become one of three cell types: polar cells, stalk cells, or main body follicle cells (Rust and Nystul, 2020). The process of pFC differentiation occurs gradually, over the course of multiple divisions and, though they are a transcriptionally heterogeneous population (Rust et al., 2020), it is not currently possible to distinguish between the pFCs in the germarium that are differentiating toward one cell type or another. Thus, here, we refer to all follicle cells in the germarium that are downstream from the FSCs as pFCs. Once Stage 1 follicles bud from the germarium, stalk cells are evident by their position between follicles, polar cells can be identified as small clusters of cells near the anterior and posterior poles of the follicles, and main body follicle cells comprise the majority of the cells around the outside of the follicle. Several markers, described below, also help identify each cell type. Outside of the germarium, mature polar cells and stalk cells are post-mitotic, whereas main body follicle cells remain proliferative until the end of Stage 6 and continue to differentiate into subtypes that correspond to the stage of oogenesis and their position on the follicle.

**Figure 1:**
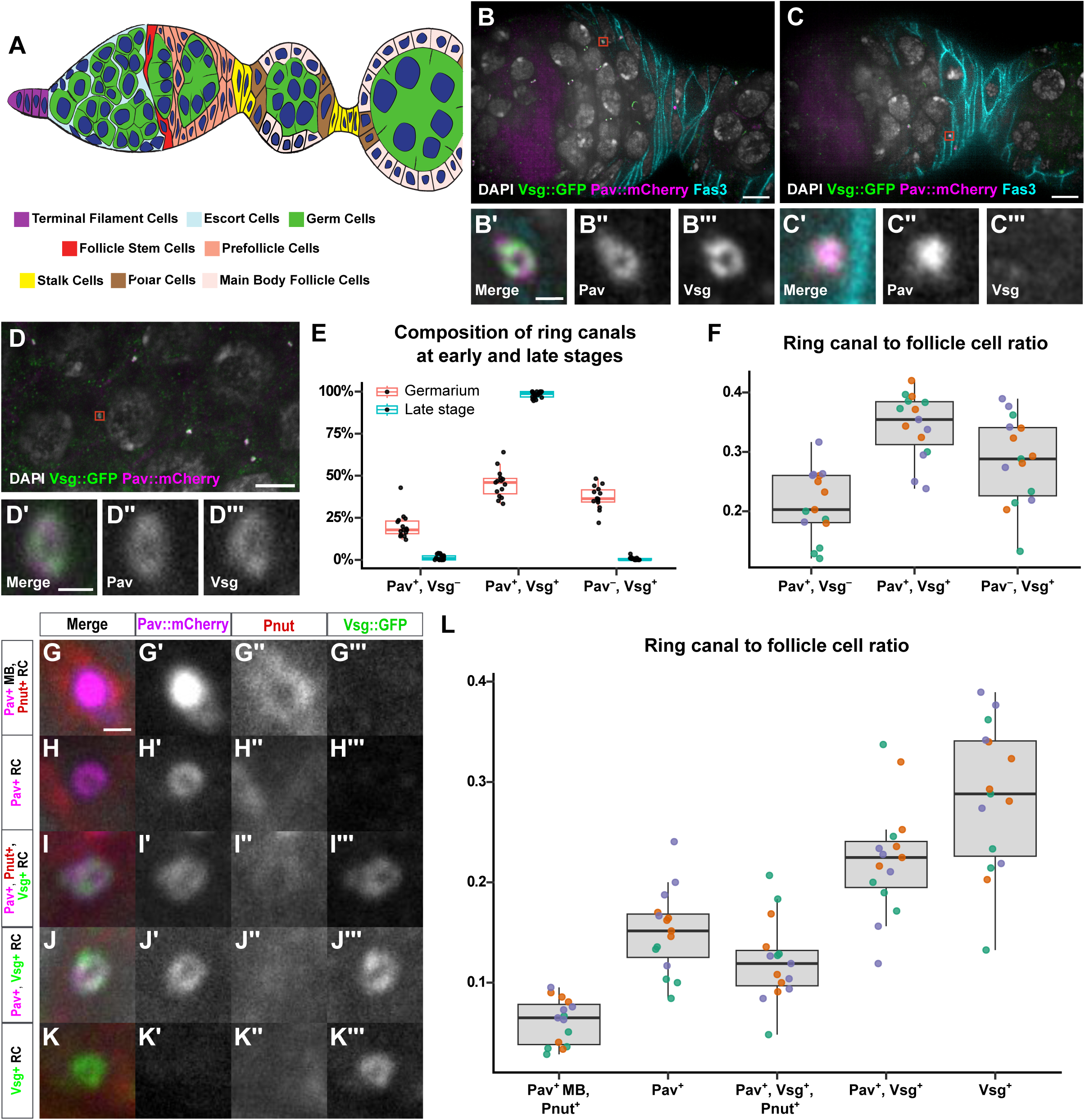
Analysis of follicle cell ring canal composition. **(A)** Illustration of the germarium and first budded follicle with the main cell types colored as indicated. **(B-C)** Ovarioles stained for DAPI (grey), Fas3 (cyan), Vsg::GFP (green), and Pav::mCherry (magenta). Pav^+^ and Vsg^+^ ring canals and Pav^+^ and Vsg^−^ midbody are outlined with red boxes (B-C) and magnified in insets (B’-B’’’ and C’-C’’’). **(D)** Optical section of late-stage follicle stained for DAPI (grey), Pav::mCherry (magenta), and Vsg::GFP (green). Co-localized Pav^+^ and Vsg^+^ ring canal are outlined with a red box (D) and magnified in insets (D’-D’’’). **(E)** Quantification of percentage of Pav^+^ and Vsg^+^ ring canal categories in the germarium and late-stage follicles. Dots are ring canal category percentage of individual germaria (n = 15) or late-stage follicles (n = 20). **(F)** Quantification of Pav^+^ and Vsg^+^ ring canal categories to total follicle cell ratio in the germarium. Dots are ring to cell ratios of individual germaria (n = 15). **(G-K)** Magnified insets of ring canals stained for DAPI (grey), Pav::mCherry (magenta), Vsg::GFP (green), and Pnut (red). Pav can form as a midbody and ring canal. Vsg and Pnut form as ring canals. **(L)** Quantification of Pav^+^, Vsg^+^, and Pnut^+^ ring canal categories in the germarium. Dots are ring to cell ratios of individual germaria (n = 15). Replicates are identified by dot color. Non-inset scale bars are 5µm. Inset scale bars are 0.5 µm.

Ring canals were first detected in *Drosophila* ovarian follicle cells by electron microscopy (Giorgi, 1978) and subsequently found to contain many of the same proteins as germline ring canals, including actin, a mucin- like protein, Visgun (Vsg), the cytokinesis proteins Cindr and Anillin, and a kinesin-like protein, Pavarotti (Pav) (Minestrini et al., 2002; Buszczak et al., 2007; Nystul and Spradling, 2007; Haglund et al., 2010; Woodruff and Tilney, 1998). Immunofluorescence imaging of Pav revealed that ring canals are absent in mature polar and stalk cells but nearly ubiquitous in main body follicle cells, with approximately 89% of main body follicle cell divisions outside of the germarium producing stable ring canals (Airoldi et al., 2011). In mature follicles, follicle cell ring canals allow passage of some individual proteins between cells (McLean and Cooley, 2013; Airoldi et al., 2011), and the observation that interconnected main body follicle cells are more likely to be in mitosis at the same time suggests that they may help coordinate proliferation (Airoldi et al., 2011). However, while Vsg^+^ and Pav^+^ ring canals have been observed in FSCs and pFCs (Nystul and Spradling, 2007; Airoldi et al., 2011), the composition and distribution of ring canals in the pFC population within the germarium, as well as the roles of ring canal proteins in pFC differentiation, are not well-understood.

Here, we apply conventional and super-resolution microscopy, genetics, and quantitative image analysis to investigate these questions. We find that ring canals in the early FSC lineage are more heterogeneous than main body follicle cell ring canals in late-stage follicles. In addition, our analysis suggests that ring canals in the early FSC lineage form through a stepwise process that begins with the elaboration of midbodies at the cytokinetic furrow and progresses through multiple stages with distinct compositions. RNAi knockdown of the cytokinesis protein Shrub (Shrb) increased the number of ring canals per cell, altered proliferation rates, and impaired pFC differentiation, whereas knockdown of the transient ring canal component, Peanut (Pnut), caused aneuploidy but did not significantly affect these other aspects of pFC biology. Taken together, these results identify ring canal formation as a point of regulation in the patterning of proliferation and differentiation in the early follicle epithelium.

## Results

### Visualizing and quantifying follicle cell ring canals in the germarium

Follicle cell ring canals are approximately 0.35 µm in diameter, which is close to the diffraction limit for conventional fluorescence microscopy. Thus, we used a spinning-disc confocal microscope with optical photon reassignment (SoRa), which provides an approximately 1.4-fold increase in resolution beyond the conventional diffraction limit (Azuma and Kei, 2015). With this system, we were able to clearly visualize follicle cell ring canals and their lumens in the germarium (**Fig. 1B**). Next, to assist in the process of determining ring canals frequency, we developed a method to efficiently quantify follicle cells in the germarium. Automated 3D segmentation of cells in the early FSC lineage is challenging because they are densely packed and have irregular, non-spherical shapes. In addition, although Fas3 is useful for distinguishing cells in the early FSC lineage from other cells in the germarium, it does not label the entire membrane and thus does not fully delineate the cell boundaries. These properties are problematic for many conventional image segmentation approaches, which use discrete global thresholds to define object boundaries and uniform shapes, such as spheres or rectangular prisms, to approximate cell morphology (Meijering, 2012; Gogoberidze and Cimini, 2024). However, a recently developed method, called StarDist, circumvents these issues by making use of higher-order star-convex polygons rather than standard bounding boxes to represent irregular cell shapes (Weigert et al., 2020). In addition, the StarDist model uses a convolutional neural network to learn boundary predictions directly from annotated training data and assigns pixels to objects by ray tracing from an object center point rather than using a global threshold. We trained StarDist to recognize follicle cells in the germarium based on Fas3 and DAPI staining, with manually created masks of follicle cell nuclei serving as ground truth. In 3D image stacks that were not used in training, the model identified follicle cells in the germarium with a precision of 69.6 ± 1.7% and a sensitivity of 48.8 ± 1.8%. This provided a useful starting point for accurately quantifying follicle cell number in each image set (**Fig. S1**).

### Follicle cell ring canals are less common and more heterogeneous in the germarium

To investigate the composition of follicle cell ring canals in the germarium, we first assessed the co-localization between Pav::mCherry and Vsg::GFP. We found both spherical Pav^+^ structures that are presumably midbodies (Echard et al., 2004) and ring-shaped Pav^+^ structures (**Fig. 1B-C**). Spherical Pav^+^ structures were always Vsg^−^, whereas 45.3 ± 8.3% of rings were Pav^+^, Vsg^+^, 37.1 ± 6.9% were Pav^−^, Vsg^+^ rings and 20.1 ± 7.5% were Pav^+^, Vsg^−^. In contrast, nearly all (98.2 ± 1.8%) of the Pav^+^ or Vsg^+^ ring canals in the post-mitotic main body follicle cells of late-stage follicles were positive for both markers (**Fig. 1D-E**). Note, we also observed some circular Pav^−^, Vsg^+^ structures in the cytoplasm of late-stage follicle cells but we did not consider them to be ring canals because they were generally larger than the Pav^+^, Vsg^+^ ring canals and not clearly localized to the plasma membrane (**Fig. S2A-C**). Because pFCs have irregular shapes and extensive intermingling of cytoplasmic projections, we cannot confidently assign each ring canal to a specific cell in the germarium. However, combining the StarDist models with manual corrections allowed us to accurately determine the total number of follicle cells in each germarium and thus to calculate the ratio of ring canal number to follicle cell number for each germarium. We found that the average ratio of Pav^+^, Vsg^−^ rings per follicle cell was 0.21 ± 0.05; of Pav^+^, Vsg^+^ rings per follicle cell was 0.34 ± 0.04; and of Pav^−^, Vsg^+^ rings per follicle cell was 0.28 ± 0.04 (n = 15) (**Fig. 1F**). Thus, consistent with previous estimates (Airoldi et al., 2011), some divisions in the early FSC lineage do not result in the formation of ring canals.

Next, we considered other common markers of ring canals, including Peanut (Pnut), phosphotyrosine (pTyr), and Hu li tai shao (Hts) (Price and Lewellyn, 2026; de Cuevas and Spradling, 1998; Robinson et al., 1994). All of these markers clearly localized to germ cell ring canals, and each exhibited a distinct localization pattern in follicle cells. Consistent with previous observations (Robinson et al., 1994), Hts localized to follicle cell membranes and partially overlapped with Pav and Vsg signals but was not noticeably enriched on ring canals compared to the adjacent regions (**Fig. S2D**). In contrast, both Pnut and pTyr were enriched on at least some Pav^+^ or Vsg^+^ rings. We frequently observed spherical Pav^+^ structures that overlapped with or were immediately adjacent to cortical Pnut signal, and many were surrounded by a clear Pnut^+^ ring (**Fig. 1G**). In addition, 34.7 ± 1.8% of Pav^+^, Vsg^+^ rings were Pnut^+^ (**Fig. S2E**). Interestingly, however, we did not observe any Pav^−^, Vsg^+^ rings that were Pnut^+^ (**Fig. 1G-L**). Conversely, the pTyr signal only colocalized with rings that were Vsg^+^, and these rings were rarely Pav^+^. Specifically, 4.91 ± 7.7% of Pav^+^, Vsg^+^ rings and 69.5 ± 10.3% of Pav^−^, Vsg^+^ rings were pTyr^+^ (**Fig. S3E-F**). Notably, structures in all of these different categories could be found throughout the early FSC lineage, including at the Region 2a/2b border, where the FSCs reside, and in both Region 2b and Region 3. Taken together, these observations indicate that ring canals are more heterogeneous in the early stages of the FSC lineage compared to later stages.

### Pnut is required for normal cytokinesis but not differentiation or ring canal formation

Pnut is a member of the septin family and a component of the contractile ring that promotes the initial ingression of the cleavage furrow (El Amine et al., 2013). Loss of *pnut* function leads to cytokinesis defects, such as binucleation and aneuploidy in multiple cell types, including follicle cells (Neufeld and Rubin, 1994; Fares et al., 1995). To knockdown *pnut* in the early FSC lineage specifically during adulthood, we combined the early follicle cell driver, *109-30-Gal4* (Hartman et al., 2010), and a temperature-sensitive *Gal80* (a combination referred to here as *109-30^ts^*) with *pnut* RNAi, and shifted adult flies to the permissive temperature for Gal4 activity (29°C) for 7 days before dissecting and analyzing the ovaries. First, we stained for Fas3 and Vasa and confirmed that RNAi knockdown of *pnut* in the early FSC lineage caused a highly penetrant aneuploidy phenotype. In 100% (n = 259) of the pnut mutant ovarioles we analyzed, the recently budded follicles contained large and misshapen cells with either a single large nucleus or two nuclei (**Fig. 2A-B**). Notably, we observed examples of aneuploid stalk, polar, and main body follicle cells, indicating that increased ploidy is compatible with differentiation toward each of these cell fates (**Fig. 2C-H**). To confirm this observation, we stained control and *pnut* mutant ovarioles for Castor (Cas), Eyes Absent (Eya), Six4, and Lamin C (LamC) as markers of distinct cell types in the early FSC lineage (Dai et al., 2017; Chang et al., 2013; Johnston et al., 2016; Anschütz et al., 2026). In wildtype ovarioles, FSCs and pFCs are Cas^+^, Eya^+^, and have a diffuse Six4 signal, whereas stalk and polar cells are Cas^+^, Eya^−^, Six4^−^, and main body follicle cells are Cas^−^, Eya^+^ with a nuclear Six4 signal. LamC is strongly enriched on the nuclear membranes of stalk cells between follicles and is largely absent from pFCs and main body follicle cells. LamC is also present on germ cell nuclear membranes and is detectable at low levels in a small number of anterior and posterior main body follicle cells. However, these other LamC^+^ cells can be distinguished from stalk cells by nuclear size, shape, and LamC signal intensity. Based on these markers, we did not detect any pFC differentiation defects in *pnut* mutant ovarioles (**Fig. S4A-D**).

**Figure 2:**
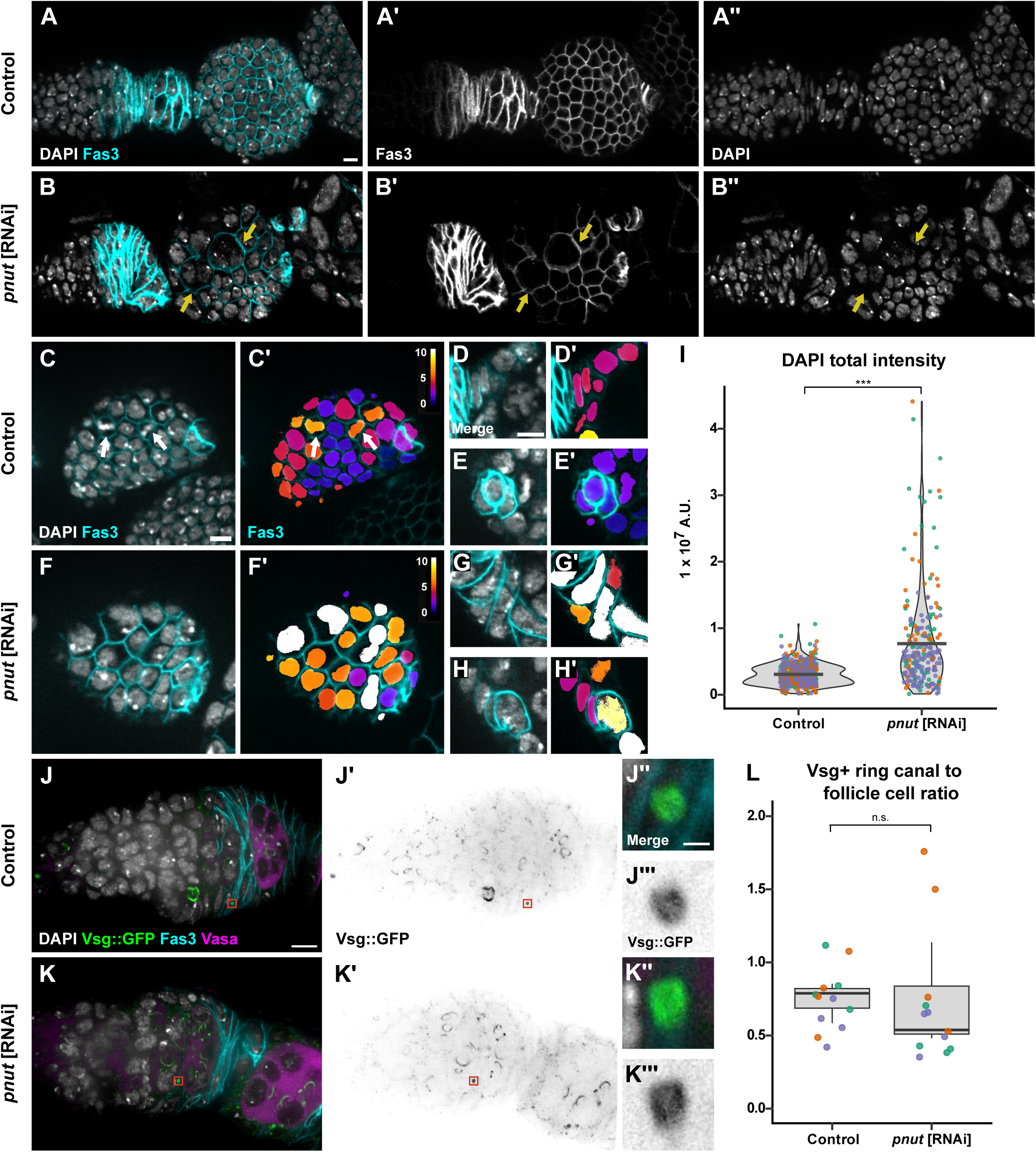
*pnut* knockdown in follicle cells causes aneuploidy but does not decrease the frequency of ring canals. **(A-H)** Ovarioles with *109-30^ts^* alone (control) or driving *pnut* [RNAi] stained for DAPI (grey) and Fas3 (cyan). Optical section of the follicle epithelium around the first budded cyst (cyan) shows typical diploid nuclear morphology in control (A) and aneuploid or binucleate nuclear morphologies in *pnut* [RNAi] (yellow arrows, B). Optical section of the follicle epithelium of a budded follicle (C, F) with insets of stalk cells, identified by their position between follicles (D, G) and polar cells, identified by the high levels of Fas3 expression (E, H). Cells in mitosis in the control, identified by their distinct shape (white arrows, C). Nuclear masks (C’-H’) pseudocolored based on total intensity of the DAPI channel within each mask. Inset scale bar is 5µm. **(I)** Quantification of follicle cell DAPI signal in follicle cells of newly budded follicles (N = 3). Dots are DAPI values of individual follicle cells. **(J-K)** Ovarioles with Vsg::GFP and *109-30^ts^* alone (control) or driving *pnut* [RNAi] stained for DAPI (grey), Fas3 (cyan), Vasa (magenta) and Vsg::GFP (green). Vsg^+^ ring canals (inverted grey) outlined with red boxes (J’-K’) and magnified in insets (J”-J’’’ and K”-K’’’). Inset scale bar is 0.5µm. **(L)** Quantification of Vsg^+^ ring canals to total follicle cell ratio in the germarium. Dots are ring to cell ratios of individual germaria (n = 12). Replicates are identified by dot color. Non-inset scale bars are 5µm. Significance values: ns= not significant, p<0.05*, p<0.01**, p<0.001*** using Welch’s t-test on average values per replicate.

To further characterize the aneuploidy phenotype, we quantified the total DAPI intensity per cell in recently budded follicles. Cells in the early stages of the FSC lineage are diploid and thus should have a ploidy ranging from 2N to 4N, depending on the phase of the cell cycle (**Fig. 2C**). Indeed, we found that the distribution of total DAPI intensities per cell in wildtype follicles was bimodal, with peaks approximately 2-fold apart (**Fig. 2I**). In contrast, the mean of the total DAPI intensities per cell in *pnut* mutant follicles was significantly higher than that of the control, and the individual values spanned a much wider range, with many cells that contained 2-4 fold more signal than even the brightest wildtype cells (**Fig. 2I**). Interestingly, there was no significant difference in the frequency of pFCs in S-phase (**Fig. S4E-G**), based on a standard EdU incorporation assay (Laws and Drummond-Barbosa, 2015). In addition, we observed many examples of *pnut* mutant aneuploid main body follicle cells in prophase or metaphase, including two adjacent cells in the same follicle (**Fig. S4H-I**), indicating that the aneuploid state does not prevent these cells from entering mitosis. Thus, consistent with the conserved function of septins to promote ingression of the cytokinetic furrow (El Amine et al., 2013), these observations strongly suggest that the aneuploidy phenotype in *pnut* mutants is caused by defects in cytokinesis. Nonetheless, despite its important role in cytokinesis, we found that *pnut* knockdown does not prevent Vsg^+^ ring canals from forming at a frequency comparable to that in controls (**Fig. 2J-L**).

### Shrb limits the number of follicle cell ring canals and is required for proper prefollicle cell differentiation

Our observation that the number of somatic ring canals in the germarium is much less than the number of pFCs (**Fig. 1G**) indicates that some pFCs undergo complete abscission. In ovarian germline stem cells, the ESCRT III protein, Shrb, promotes abscission by localizing to the cytokinetic furrow and destabilizing ring canals (Mathieu et al., 2022). To investigate whether Shrb has a similar function in the early FSC lineage, we combined Vsg::GFP with *109-30^ts^* and *shrb* RNAi and quantified the ratio of ring canals to pFCs after 7 days at the permissive temperature for Gal4 activity. Indeed, we found that the average ratio of Vsg^+^ rings per pFC was significantly higher in *shrb* mutant germaria compared to the control (0.97 ± 0.1 vs. 0.75 ± 0.1, respectively, N = 3, p < 0.05) (**Fig. 3A-C**). This indicates that Shrb inhibits ring canal formation or stability in the early FSC lineage.

**Figure 3:**
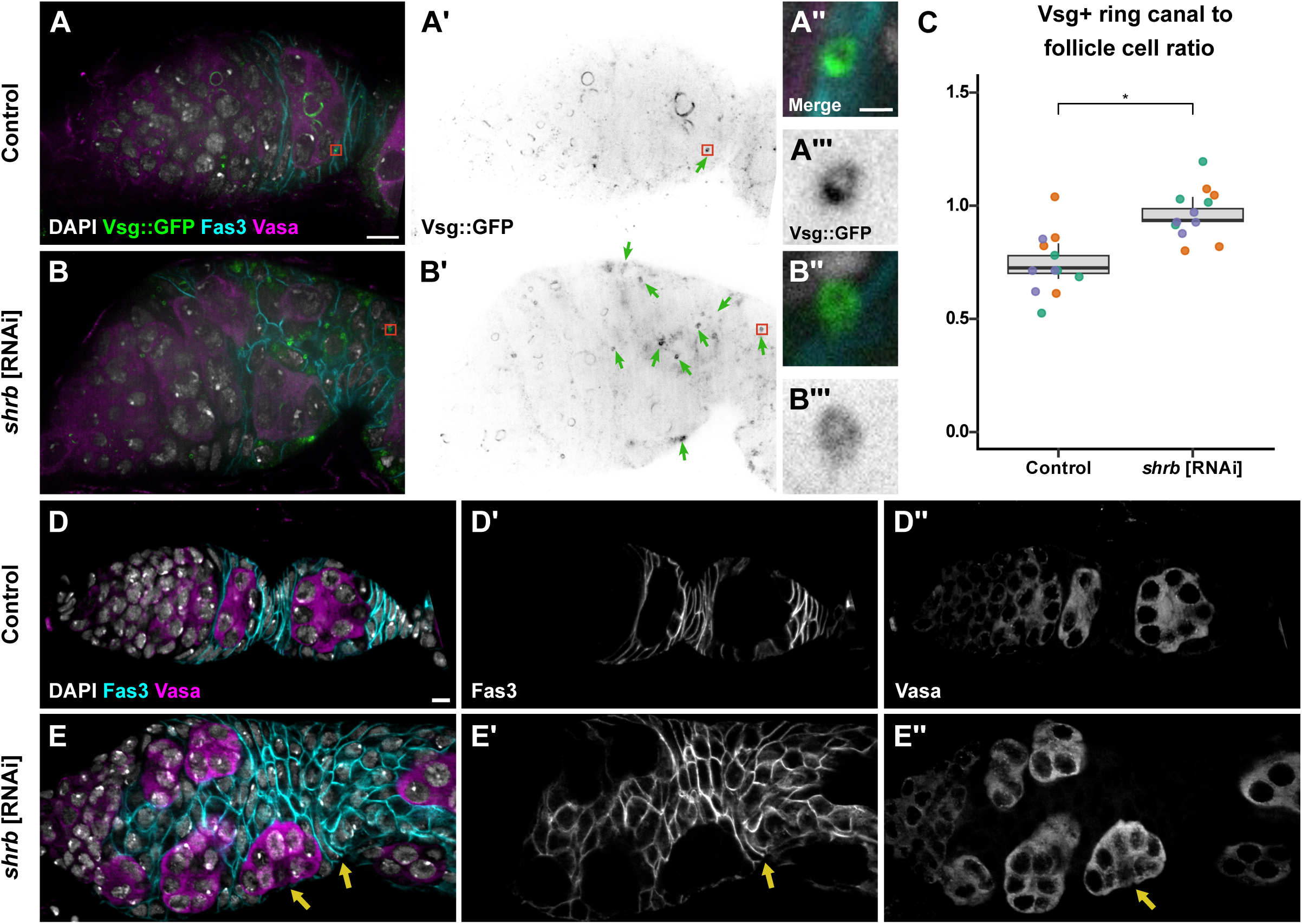
*shrb* knockdown in follicle cells disrupts tissue patterning and increases the frequency of ring canals. **(A–B)** Ovarioles with Vsg::GFP and *109-30^ts^* alone (control) or driving *shrb* [RNAi] stained for DAPI (grey), Fas3 (cyan), Vasa (magenta) and GFP (green). Vsg^+^ ring canals (inverted grey) in control and *shrb* [RNAi] germaria, outlined with red boxes and green arrows (A’-B’) and magnified in insets (A”-A’’’ and B’’-B’’’)**. (C)** Quantification of Vsg+ ring canals to total follicle cell ratio in the germarium. Dots are ring to cell ratios of individual germaria (n = 12). **(D-E)** Ovarioles stained for DAPI (grey), Fas3 (cyan) and Vasa (magenta) have a typical morphology in control (D) and a disorganized follicle epithelium and impaired budding of germ cell cysts with *shrb* [RNAi] (yellow arrows, E). Replicates are identified by dot color. Non-inset scale bars are 5µm. Inset scale bar is 0.5µm. Significance values: ns= not significant, p<0.05*, p<0.01**, p<0.001*** using Welch’s t-test on average values per replicate.

To investigate whether knockdown of *shrb* also disrupts the patterning of proliferation or differentiation in the early FSC lineage, we stained control and *shrb* mutant ovarioles for Fas3 and Vasa. We found that *shrb* knockdown caused a strong, highly penetrant phenotype (100%, n = 180) characterized by an increased number of follicle cells in the germarium and severe budding defects. Specifically, follicles accumulated in a disorganized pattern at the posterior edge of the germarium, rather than budding off in a single file, and the cells between follicles failed to form into a single row of stalk cells (**Fig. 3D-E**). In addition to promoting cytokinesis, ESCRT complexes have other functions in the cell, including the recycling of cell surface receptors and endosomal sorting (Park et al., 2024; Vaccari et al., 2009; Stoten and Carlton, 2018). In *Drosophila* ovarioles, germ cells that are mutant for *Vps28* or *Vps25* (ESCRT I and II genes, respectively) and follicle cells mutant for *Vps22* (an ESCRT II gene) show defects in the cortical actin cytoskeleton (Vaccari et al., 2009). However, *109-30^ts^* driven expression of RNAi constructs against the ESCRT I complex protein, TSG101, or the ESCRT II complex protein, Vps25, with *109-30^ts^* did not phenocopy the *shrb* RNAi phenotype in the follicle epithelium (**Fig. S5A-E**), suggesting that this phenotype is not due to a general loss of ESCRT complex functions.

To further investigate the phenotypes caused by *shrb* knockdown, we stained for markers of pFCs and stalk cells. We found that nearly all *shrb* mutant ovarioles had an increased number of cells that expressed Cas and LamC distributed throughout Regions 2b and 3 of the germarium and in the stalk cell regions (**Fig. 4A-D**). In contrast, we did not observe any LamC^+^ pFCs in Region 2b in control germaria and, the rare LamC^+^ cells we observed in Region 3 typically had lower LamC signal and were located near the region where a new stalk was forming (**Fig. S5F**). Lastly, we performed an EdU incorporation assay to measure proliferation rates. Using the StarDist pipeline to quantify the number of follicle cells in each germarium, we found a significant increase in the average percent of EdU^+^ cells in *shrb* mutants compared to the control (23.9 ± 4.0% vs 11.7 ± 2.7%, respectively) (**Fig. 4E-G)**. Since *shrb* mutant ovarioles have more LamC^+^ cells, we hypothesized that at least some of this increase in EdU^+^ cells may be due to increased proliferation in pFCs that are differentiating toward the stalk cell fate. To test this idea, we co-stained control and *shrb* mutant germaria for LamC and EdU. LamC^+^, EdU^+^ cells were rare in control germaria (0.9 ± 1.3 cells per germarium, n = 30), as expected, since mature stalk cells are postmitotic. In contrast, we observed significantly more LamC^+^, EdU^+^ cells in *shrb* mutant germaria (18.2 ± 3.3 cells per germarium, n = 28) (**Fig 4H-J)**. In summary, Shrb is required to inhibit ring canal formation or stability in a subset of pFCs, reduce the overall rate of proliferation in the pFC population, and facilitate proper patterning of the stalk cell identity.

**Figure 4:**
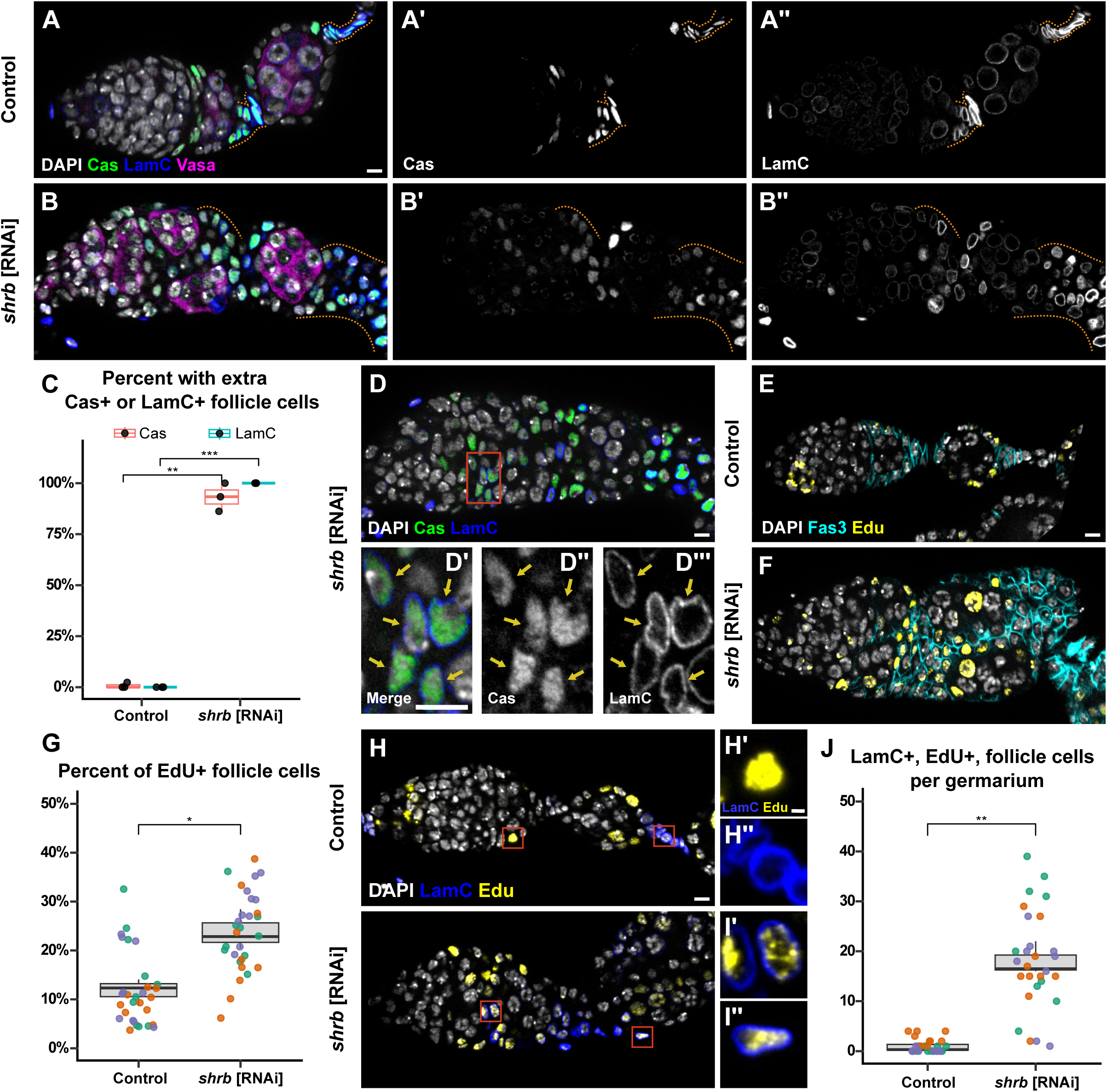
*shrb* knockdown in follicle cells increases proliferation and disrupts differentiation. **(A-B)** Ovarioles with *109-30^ts^* alone (control) or driving *shrb* [RNAi] stained DAPI (grey), Cas (green), LamC (blue), and Vasa (magenta). The stalk cell region (orange highlighted dotted lines) contains an increased number of LamC^+^ cells in the *shrb* [RNAi] group compared to control. **(C)** Quantification of the number of germaria with excess Cas^+^ or LamC^+^ cells in control (n = 121) and *shrb* [RNAi] (n = 131). Dots are replicate means of germaria with an excess number of cells that are positive for each marker. **(D)** Ovariole with *109-30^ts^* driving *shrb* [RNAi] stained for DAPI (grey), Cas (green) and LamC (blue). Cas^+^ and LamC^+^ cells in Region 2b in *shrb* [RNAi] germarium, outlined with red box (D) and magnified in insets (D’-D’’’). Yellow arrows indicate pFCs. Inset scale bar is 5 µm. **(E-F)** Ovarioles stained for DAPI (grey), Fas3 (cyan), and EdU (yellow). **(G)** Quantification of the percent of EdU^+^ follicle cells in the germarium. Dots are EdU to cell ratios of individual germaria (n = 30). **(H-I)** Ovarioles stained for DAPI (grey), LamC (blue), and EdU (yellow). Magnified in insets of Edu^+^ and LamC^−^ or LamC^+^ and EdU^−^ cells in control (H’-H’’) or LamC^+^ and EdU^+^ cells in *shrb* [RNAi] (I’-I’’). Inset scale bar is 1µm. **(J)** Quantification of the number of LamC^+^ and EdU^+^ cells per germarium. Dots are LamC+ cells of individual germaria (n = 28-30). Non-inset scale bars are 5µm. Significance values: ns= not significant, p<0.05*, p<0.01**, p<0.001*** using Welch’s t-test on average values per replicate.

## Discussion

Our studies have provided several new insights into the composition and function of ring canals in the early FSC lineage. First, our findings support a model in which ring canals in prefollicle cells progress through a series of distinct molecular states. In multiple species, germ cell ring canals are known to derive from Pav/MKLP1^+^ midbodies, whereas the somatic cells of the Drosophila testis do not appear to undergo a similar midbody-to-ring canal transition (Price et al., 2023; Greenbaum et al., 2007). Here, we frequently observed spherical Pav^+^ structures surrounded by Pnut, suggesting that follicle cell ring canal formation begins with recruitment of Pnut to a Pav^+^ midbody followed by the elaboration of the midbody into a Pav^+^ ring. However, we cannot confirm this with live imaging because mCherry and GFP are quickly bleached by SoRa super-resolution imaging. Next, Pnut is likely lost as Vsg is recruited, accounting for the intermediate Pav^+^ and Pav^+^, Vsg^+^, Pnut^+^ structures. The Pav^+^, Vsg^+^ state is likely to be a mature state since this composition predominates at later stages of the follicle cell lineage and Pav^+^ follicle cell ring canals are capable of facilitating intercellular transport (Airoldi et al., 2011; McLean and Cooley, 2013). Thus, our observations support a general progression from an early Pav^+^, Vsg^−^ state to a mature Pav^+^, Vsg^+^ state, with Pnut transiently associated with at least some intermediates. The Pav^−^, Vsg^+^ rings may be an alternative form, or they may be in the process of breaking down.

Although pTyr is one of the earliest known markers of germ cell ring canals (Robinson and Cooley, 1997; Robinson et al., 1994), this study provides the first evidence to our knowledge of a pTyr signal on follicle cell ring canals. In germ cell ring canals, tyrosine phosphorylation of the actin-binding protein Kelch is induced by the Src64/tec29 kinase cascade and contributes to the regulation of ring canal growth and stability in late oogenesis (Kelso et al., 2002; Roulier et al., 1998; Guarnieri et al., 1998; Dodson et al., 1998). However, Kelch is not present in follicle cell ring canals, and Src64 is not required in pFCs for proper proliferation or differentiation (O’Reilly et al., 2006). Thus, although the kinase cascade and its targets are different in follicle cell ring canals, our analysis suggests that the phosphorylation of one or more pFC ring canal components may facilitate the transition from the Pav^+^, Vsg^+^ state to the Pav^−^, Vsg^+^ state. In germ cells, the sequential recruitment of ring canal proteins spans across multiple stages of differentiation, with phosphotyrosine signal detectible in Region 1 but the recruitment of Hts and actin delayed until after germ cells exit mitosis in Region 2a. In contrast, we did not find any category that was absent from the earliest stages of the FSC lineage, suggesting that pFCs progress through these categories over a relatively short time span as they move through the germarium.

Second, our findings demonstrate that the decision to undergo complete abscission or maintain a stable ring canal is a key point of regulation in pFC differentiation (**Fig. 5**). In support of this, we confirmed the prediction that some but not all pFC divisions in wildtype ovarioles produce stable ring canals (Airoldi et al., 2011). The variation in both the presence and the composition of ring canals that we observed is likely a reflection of the underlying heterogeneity in the developmental trajectories of the different pFCs in the germarium. In addition, we found that knockdown of *shrb* stabilizes ring canals in the germarium, increasing the ratio of ring canals to cells to approximately 100%, indicating that essentially all divisions resulted in a stable ring canal. Given the proliferation and differentiation phenotypes we observed in *shrb* mutants, this strongly suggests that the dissolution of ring canals is required for pFCs to differentiate toward the postmitotic stalk cell fate. This may be necessary to prevent these pFCs from continuing to receive cell cycle cues from sister cells that are still in the cell cycle. It seems unlikely that the premature expression of LamC^+^ in some pFCs in *shrb* mutant germaria is also due to aberrant transfer of cytoplasmic contents since this marker is normally expressed much later. However, the overall tissue architecture is severely disrupted in *shrb* mutants, so it is possible that this phenotype results from mislocalization of mature stalk cells or from a disruption in the patterning of stalk cell cues.

**Figure 5:**
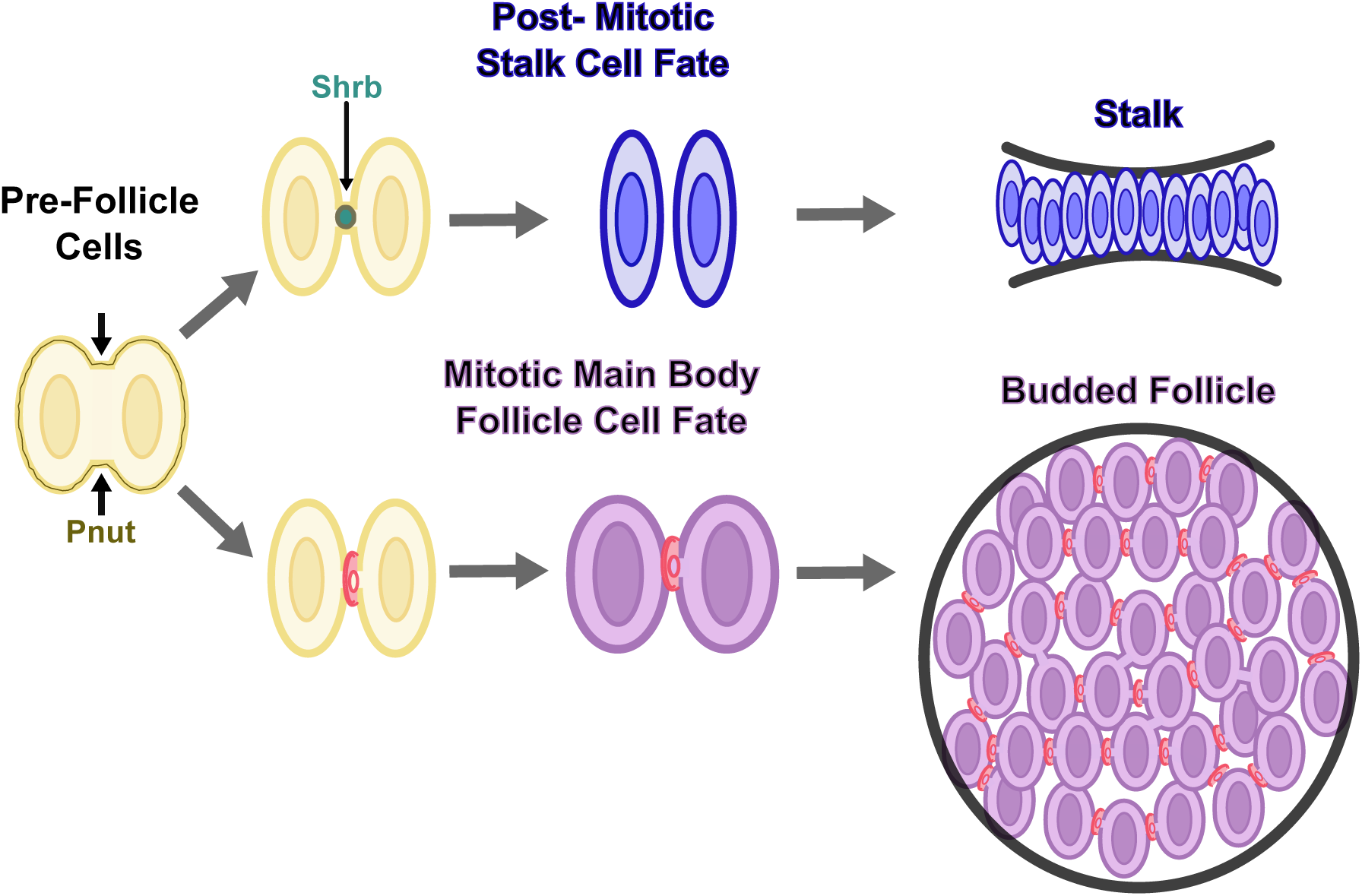
Model for the modes of cytokinesis in prefollicle cells. Illustration depicting different modes of cytokinesis along different cell fate trajectories. During cytokinesis in pFCs, Pnut aids in the constriction of the cleavage furrow. Following cleavage furrow ingression, pFCs can undergo either incomplete or complete cytokinesis. Cells that progress toward a post-mitotic stalk cell fate complete abscission through a process that requires Shrub. This action allows the separation of stalk cells, facilitating their organization into a single row. Alternatively, in cells that progress toward a mitotic main body follicle cell fate, the cleavage furrow arrests and a ring canal structure is formed. Ring canals in main-body follicle cells are abundant in a budded follicle, which facilitates intercellular communication that may be important for coordinated behaviors.

Building on these observations, we found that, although knockdown of *pnut* caused a highly penetrant aneuploidy phenotype, it did not significantly alter the ratio of ring canals per follicle cell or interfere with the specification of the major cell types in the early FSC lineage. The presence of ring canals in this mutant background indicates that the formation and maintenance of follicle cell ring canals do not require a fully functional contractile ring. This implies that, even in this mutant context, these pFCs retain the ability to either complete abscission or form a ring canal as needed to acquire the appropriate cell fates. Understanding the functional significance of the different ring canal compositions in pFCs and how they are both regulated by and contribute to cell fate decisions in pFCs will be an important topic for further investigation.

Lastly, an additional advance provided by this study is the development of a StarDist model that automates the segmentation of follicle cell nuclei in the germarium. This model is an excellent starting point for accurately quantifying the number of follicle cells per germarium. A relatively straightforward process of visually inspecting the results to count false positives and false negatives can be applied to further increase accuracy. In addition, even without manual corrections, this model may be useful for efficiently extracting quantitative information, such as position and signal intensity, from a large population of cells that is highly enriched for follicle cells in the germarium. Importantly, the accuracy of these models can be further improved in the future through a bootstrapping process in which the model is applied to a novel set of images and manual corrections are used to create new ground truth data for additional training. In this way, the models can continue to evolve with use.

The discovery of intercellular bridges across many different somatic cell types from multiple species demonstrates their widespread conservation. Though there are likely substantial differences in how these structures form and function in these diverse contexts, much can be learned from studying specific examples in detail. Future studies of follicle cell ring canals will help elucidate the upstream cues that pattern the formation and maintenance of these fascinating structures in the pFC population and the downstream effectors of proliferation and differentiation that depend on them.

## Materials and Methods

### Fly Husbandry

All stocks were maintained on standard growth media (6% cornmeal, 5% molasses, 1.2% ethanol, 1.1% yeast, 0.6% agar, 0.4% propionic acid, 0.2% tegosept in deionized water). Parent fly stocks and non-temperature-sensitive crosses were maintained at 25 °C. Temperature-sensitive crosses were kept at 18 °C until the F1 progeny eclosed. F1 progeny were shifted to 29 °C for 1 week and fed wet yeast prior to dissections.

### Fly Stocks

UAS driven constructs were combined with *109-30-Gal4* and *tub-Gal80^ts^*. *109-30-Gal4* is expressed in the early FSC lineage and Gal80^ts^ restricts Gal4 activity at 18 °C, but not 29 °C. Fly stocks used and their accompanying information are listed below.

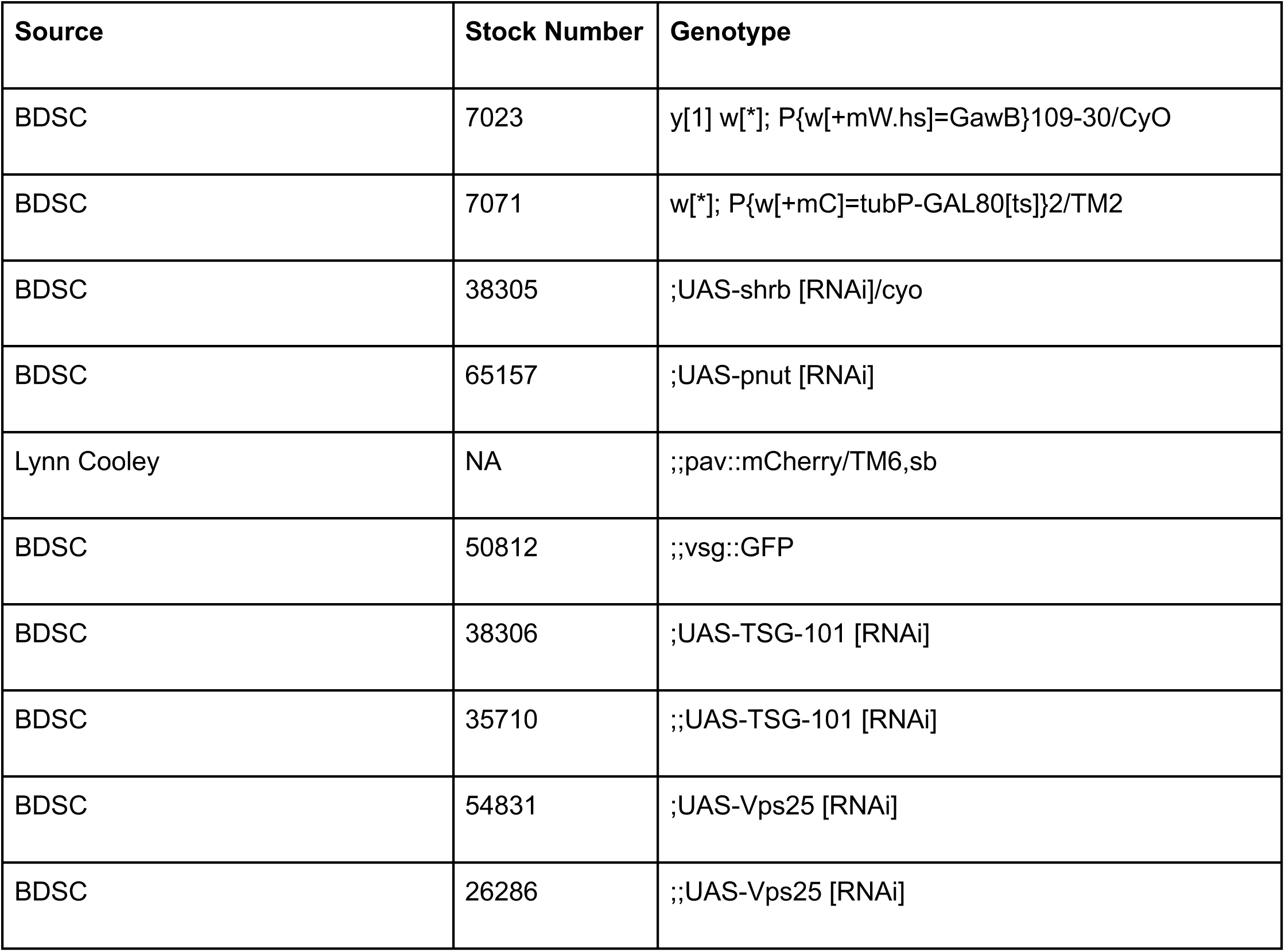

### Immunofluorescence

Drosophila ovaries were dissected in 1x PBS media and subsequently fixed in 4% PFA + PBS for 20 minutes. Following fixation, the ovaries were washed twice with 1x PBS and prepped for antibody incubation with 1X PBS + 0.2 Triton X-100 0.2% + 0.5% bovine serum albumin (block) twice. Primary antibodies were diluted in block solution and incubated with the samples overnight on a nutator at 4 °C in the dark. Then, the samples were washed with block twice, and incubated in block for an hour on a nutator at 25 °C in the dark. Secondary antibodies were diluted in block solution and incubated with the samples overnight on a nutator at 4 °C in the dark. Then, the samples were washed twice with block and 1X PBS and mounted on glass slides in DAPI-fluoromount G Slide Mounting Media. The information about the primary and secondary antibodies used in this study is listed below.

### Primary Antibodies

| Source | Species | Antigen | Concentration (µL) |
| --- | --- | --- | --- |
| Developmental Studies Hybridoma Bank | Mouse | Fascillin-III | 1:100 |
| Developmental Studies Hybridoma Bank | Mouse | Peanut | 1:100 |
| Developmental Studies Hybridoma Bank | Mouse | Eya | 1:50 |
| Developmental Studies Hybridoma Bank | Mouse | LaminC | 1:100 |
| Developmental Studies Hybridoma Bank | Mouse | Hu-li tai shao | 1:50 |
| Developmental Studies Hybridoma Bank | Rat | Vasa | 1:500 |
| Chromotek | Rat | RFP | 1:1000 |
| Torrey Pines | Rabbit | GFP | 1:1000 |
| Abcam | Rabbit | Phospho-tyrosine | 1:1000 |
| A gift from Jonathan Benito Sipos | Rabbit | Castor | 1:500 |
| A gift from Eric Rulifson | Rabbit | Six4 | 1:1000 |

### Secondary Antibodies

| Company | Species | Wavelength | Concentration (µL) |
| --- | --- | --- | --- |
| Invitrogen | Goat-Mouse | 488 | 1:1000 |
| Invitrogen | Goat-Mouse | 555 | 1:1000 |
| Invitrogen | Goat-Mouse | 647 | 1:1000 |
| Invitrogen | Goat-Rat | 488 | 1:1000 (1:500 for primary antibody, vasa) |
| Invitrogen | Goat-Rat | 555 | 1:1000 (1:500 for primary antibody, vasa) |
| Invitrogen | Goat-Rat | 647 | 1:1000 (1:500 for primary antibody, vasa) |
| Invitrogen | Goat-Rabbit | 488 | 1:1000 |
| Invitrogen | Goat-Rabbit | 555 | 1:1000 |
| Invitrogen | Goat-Rabbit | 647 | 1:1000 |

### EdU staining and quantification

Drosophila ovaries were dissected and incubated with EdU solution (1µl EdU and 499µl 1x PBS) from the Click-iT Alexa Fluor 555 Imaging Kit for one hour. The samples were then washed with 1X PBS twice and fixed in 4% PFA + 1X PBS for 10 minutes. Next, the samples were washed twice with 1x PBS, then placed in 1X PBS + 0.2% Triton X-100 + 0.5% bovine serum albumin (block) for 1 hour at 25 °C. Primary antibody dilution concentrations were prepared in block and incubated overnight on a nutator at 4 °C in the dark. Next, the samples were washed twice with block and incubated in block on a nutator for one hour at 25 °C in the dark. Then, the samples were washed with 1X PBS twice and incubated in a Click-iT reaction mixture (430µl 1X Click-iT Buffer, 20µL CuSO4, 1.2µL Alexa Fluor, and 50 µL Reaction Buffer Additive) for one hour on a nutator at 25 °C in the dark. The Click-iT reaction mixture was washed from ovary samples with 1X PBS four times and the samples were incubated in block for one hour on a nutator at 25 °C in the dark. The secondary antibodies were diluted in block and incubated overnight on a nutator at 4 °C in the dark. Then, ovaries were washed twice and mounted on glass slides in DAPI-fluoromount G Slide Mounting Media. To quantify the number of EdU^+^ cells per follicle cell in control and *shrb* [RNAi] samples, the total follicle cells were quantified using the Stardist model and the EdU^+^ follicle cells were identified based on detection of EdU signal above a set threshold. For remaining EdU analysis, EdU^+^ follicle cells were manually quantified.

### Imaging

Samples were imaged using a Zeiss M2 AxioImager with an Apotome unit and a 63x lens, or a Nikon SoRa. In all experiments in which image intensity was quantified, the images of control and experimental samples were acquired using the same microscope settings, including exposure times and light intensities. Image processing was done using FIJI, keeping brightness and contrast settings constant when appropriate, and using a standard pipeline to process raw images into figure panels. Images were pseudocolored using the Cyan, Magenta, Grays, Green, Red, or Fire lookup tables available in FIJI. The three-dimensional reconstructions of ring canal images were created using Imaris v9.3.1.

### Ring canal quantification

Ring canals were identified by the presence of a lumen and the resemblance of a ring structure. All ring canal counts were gathered from image stacks of fixed Drosophila ovarioles. Ring canal structures were visualized by staining for Vsg::GFP, Pav::mCherry, and/or Pnut. Midbody structures were identified as Pav^+^ spherical structures.

### StarDist training and performance evaluation

A StarDist 3D model was trained on 36 image stacks of wildtype germaria and 8 image stacks of *shrb* RNAi germaria stained for Fas3 and DAPI with manually created masks of the nuclei of Fas3^+^ cells as ground truth. A transfer learning approach was performed, using the StarDist 3D_demo model as a starting point. The images were split into training and validation sets, and the model underwent 400 training epochs. Data augmentation techniques were employed using the StarDist augmenter function. StarDist model performance was evaluated by comparing predicted 3D instance labels with manually curated counts on images of wildtype germaria stained for Fas3 and DAPI that were not part of the training dataset.

Image segmentation to quantify the number of follicle cells in a set of image stacks was performed by applying the StarDist segmentation pipeline to each image stack, visualizing the results in FIJI, and using the counter tool to count the number of masks on Fas3^−^ cells and duplicate masks on Fas3^+^ cells (false positives) and Fas3^+^ cells without masks (false negatives). For analysis of ploidy in the control and *pnut* mutant images, masks were manually refined to more accurately delineate the regions in each image stack that contained DAPI signal.

### Statistics and Data Availability

Tests for statistical significance were calculated using the statistical tools in R. Graphs were created in R using tidyverse packages. Panels for figures were assembled in Inkscape. No data points from the experiments shown in the figures were excluded. Samples were not randomized or blinded. Raw data for all the quantifications and all code used for image segmentation and data analysis are available at https://github.com/NystulLab/MendozaRingCanal.git.

## Acknowledgements

We are grateful to the Bloomington Drosophila Stock Center and the Developmental Studies Hybridoma Bank for stocks and reagents, and Lynn Cooley for stocks. We are thankful for the well-curated and very helpful resource, FlyBase. We thank Anders Nystul for help with manual refinement of the masks used for the analysis of ploidy in control and *pnut* mutant image stacks. Lastly, we thank Emily Wolfgram, Rhiannon Roarke, and Sarah Soliman for helpful discussions and feedback.

## Figure Legends

**Supplemental Figure 1. Training and evaluation of the StarDist model.**

To train the StarDist model, we (1) acquired 3D confocal image stacks of ovarioles stained for DAPI (gray) and Fas3 (cyan); (2) manually annotated follicle cell nuclei using the image annotation tools in napari; and (3) converted the images to 8-bit, resized the images and masks to approximately 250 pixels in the x and y dimensions, and trained the StarDist model using the pipeline provided in the StarDist GitHub repository.

To apply the trained model, we (4) acquired independent 3D confocal image stacks, rotated and cropped them to include the follicle cell region of the germarium, and processed them as described for the training images; and (5) applied the trained StarDist model to segment follicle cell nuclei.

To evaluate model performance, predicted nuclear masks were manually scored in a subset of images as true positives (TP), false positives (FP), or false negatives (FN). A true positive was defined as a predicted mask corresponding to a follicle cell nucleus, a false positive as a predicted mask that did not correspond to a follicle cell nucleus, and a false negative as a follicle cell nucleus that was not detected by the model. Boxplots show precision and recall, calculated as follows:

Precision = TP / (TP + FP)

Recall = TP / (TP + FN)

**Supplemental Figure 2: Additional analysis of follicle cell ring canal composition**

**(A-C)** Optical section of late-stage follicle with Pav::mCherry and Vsg::GFP stained for DAPI (grey), Pav::mCherry (magenta), and Vsg::GFP (green). Merge 1 and merge 2 ring structures are outlined with red boxes (A) and magnified insets (B-C) in a late-stage follicle. **(B-C)** Merge 1 depicts a standard ring canal size and shape that is Pav^+^ and Vsg^+^, while merge 2 displays an abnormal ring-like size and shape that is Pav^−^ and Vsg^+^. The Pav^−^ and Vsg^+^ ring-like structure does not localize to the membrane and possess similar qualities as typical follicle cell ring canals. **(D)** Ovariole stained for DAPI (grey), Pav::mCherry (magenta), Vsg::GFP (green), and Hts (red). Multiple Pav^+^ and Vsg^+^ ring canals, outlined with blue box (D) and magnified in insets (D’-D’’’’). Hts does not form a ring-shaped structure. **(E)** Quantification of the percentage of ring canals that are Pnut^+^ among all Pav^+^ and Vsg^+^ ring canals in the germarium. Dots are Pnut^+^ ring canals among other ring canal categories in individual germaria (n = 15). Replicates are identified by dot color. Non-inset scale bars are 5µm. Inset scale bars are 0.5µm.

**Supplemental Figure 3: Analysis of phosphotyrosine on follicle cell ring canals**

**(A)** Ovariole with Pav::mCherry and Vsg::GFP stained for DAPI (grey), Pav::mCherry (magenta), Vsg::GFP (green), and pTyr (red). Merge 1, merge 2, and merge 3 ring canals are outlined with blue boxes (A) and magnified insets (B-D) in the germarium. **(B-D)** Merge 1 depicts strong pTyr signal surrounding a Pav^−^ and Vsg^+^ ring canal. Merge 2 shows little to no pTyr signal around a Pav^+^ and Vsg^+^ ring canal. Merge 3 demonstrates no pTyr signal around a Pav^+^ and Vsg^−^ ring canal. **(E)** Quantification of pTyr ^+^ and pTyr^−^ signal co-**localized** with various ring canal categories, separated by color, in multiple replicates of individual germaria (N = 3). **(F)** Quantification of the percent of ring canal categories that are also pTyr^+^. Dots are the replicate means of pTyr^+^ ring canals among all ring canal categories in individual germaria (N = 3). Replicates are identified by dot color. Non-inset scale bars are 5µm. Inset scale bars are 0.5µm.

**Supplemental Figure 4: *pnut* knockdown in follicle cells causes aneuploidy but does not impair prefollicle cell differentiation**

**(A-D)** Optical section of the follicle epithelium with *109-30^ts^*alone (control) or driving *pnut* [RNAi] stained for DAPI (grey) and either Six4 (green) and Eya (magenta) (A-B) or Cas (green) and LamC (blue) (C-D). **(E-F)** Ovarioles stained for DAPI (grey), Fas3 (cyan) and EdU (yellow). **(G)** Quantification of the percentage of EdU^+^ cells in control and *pnut* [RNAi] germaria. Dots are EdU to cell ratios of individual germaria (n = 30). Replicates are identified by dot color. **(H-I)** *pnut* [RNAi] ovariole stained for DAPI (grey) and Fas3 (cyan), showing adjacent aneuploid cells in prophase and metaphase, respectively (yellow arrows). Significance values: ns= not significant, p<0.05*, p<0.01**, p<0.001*** using Welch’s t-test on average values per replicate. Scale bars are 5µm.

**Supplemental Figure 5: Knockdown of ESCRT I and ESCRT II genes in follicle cells does not impair prefollicle cell differentiation**

**(A-E)** Ovarioles with *109-30^ts^* alone (control) or driving multiple RNAi lines of ESCRT family genes stained for DAPI (grey), Fas3 (cyan), and Vasa (magenta). No severe phenotypes observed in the ESCRT I gene, *TSG101* [RNAi] (B-C) and ESCRT II gene, *Vps25* [RNAi] (D-E). **(F)** Control ovariole stained for DAPI (grey) and LamC (blue). Low LamC+ signal can be observed near stalk cell region (yellow arrows) in the germarium (F’). Scale bars are 5µm.

