## Supplemental files 1-5 for "Cytokinesis genes regulate ring canal patterning and follicle cell differentiation in *Drosophila*"

**Supplemental Figure 1**

##### 1. Acquire 3D confocal stacks

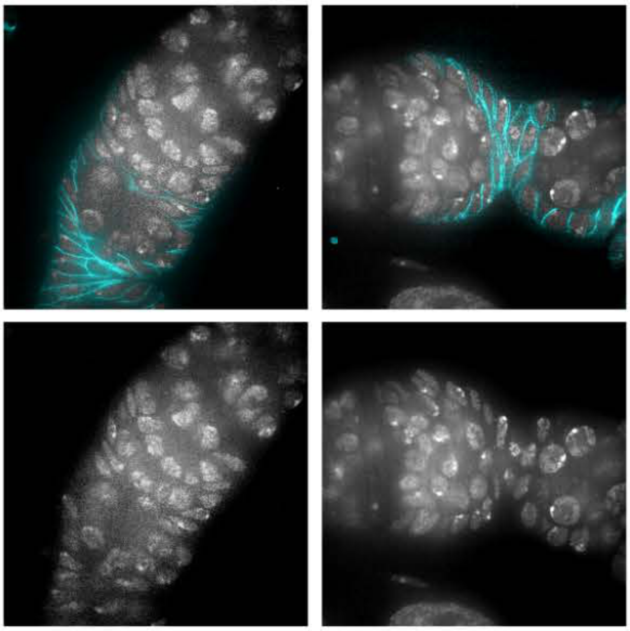

##### 2. Manually create FC masks

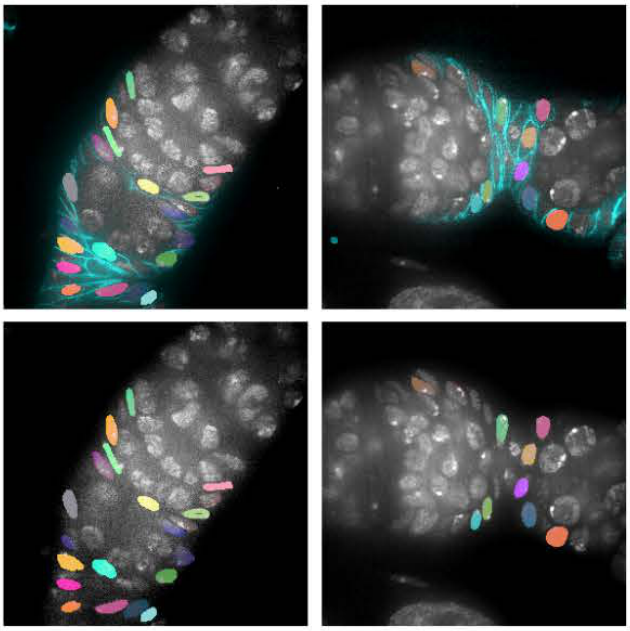

##### 3. Train Stardist on resized images with manually created masks

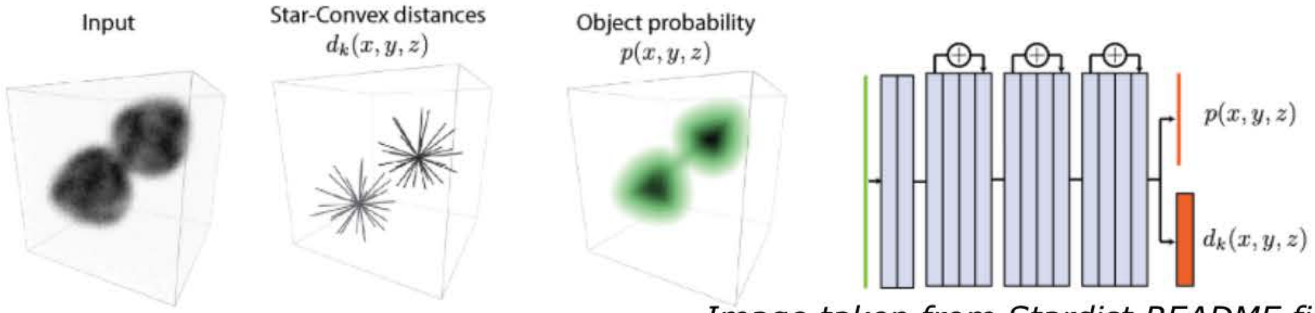


*Image taken from Stardist README file*

##### 4. Acquire new 3D confocal stacks, rotate, crop and resize

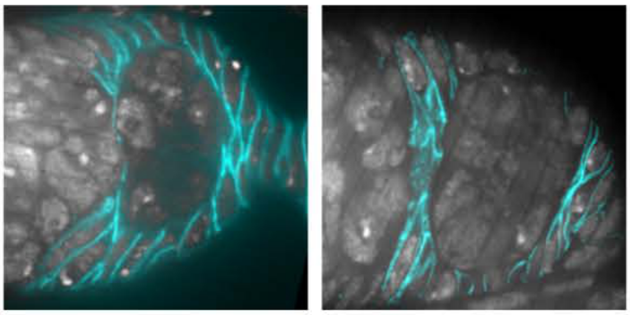

##### 5. Apply Stardist model to predict FC nuclei

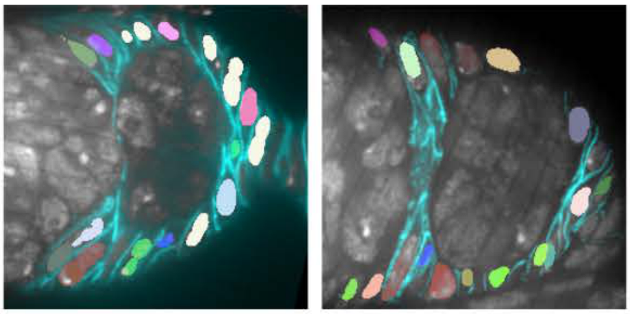

##### Comparison of Stardist model to maually generated ground truth

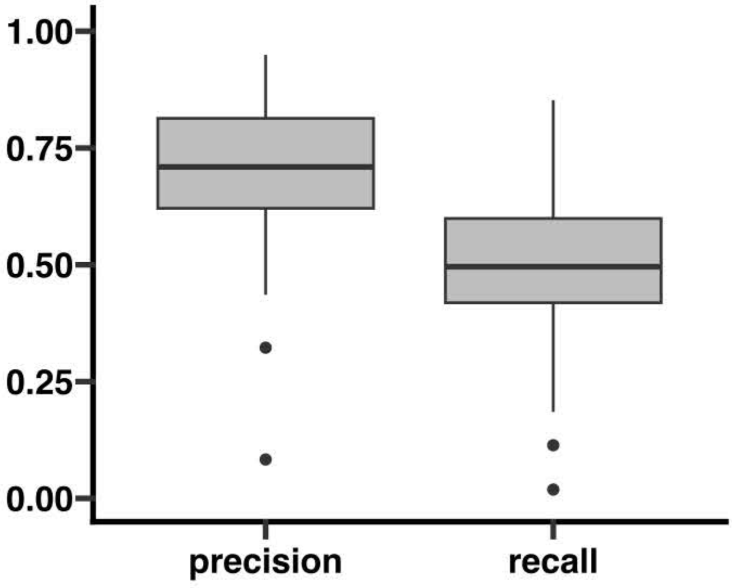

### Supplemental Figure 2

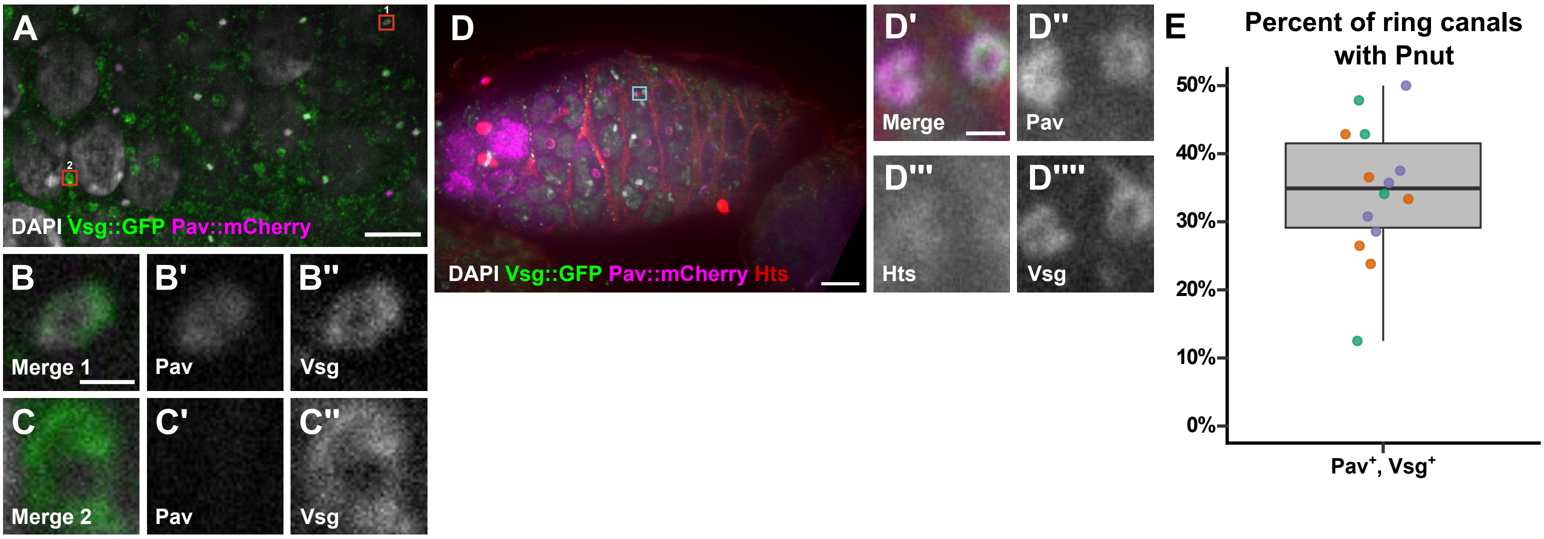

### Supplemental Figure 3

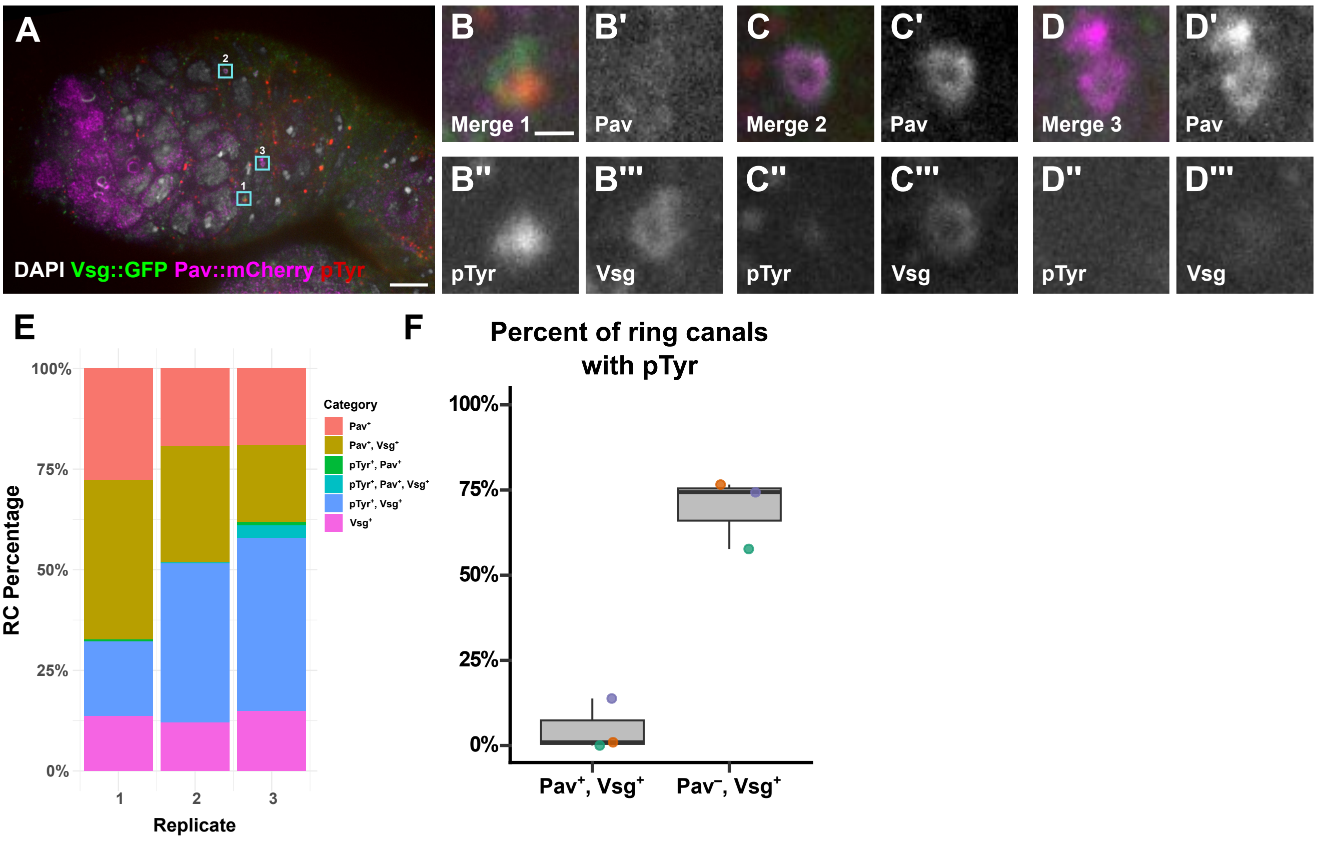

### Supplemental Figure 4

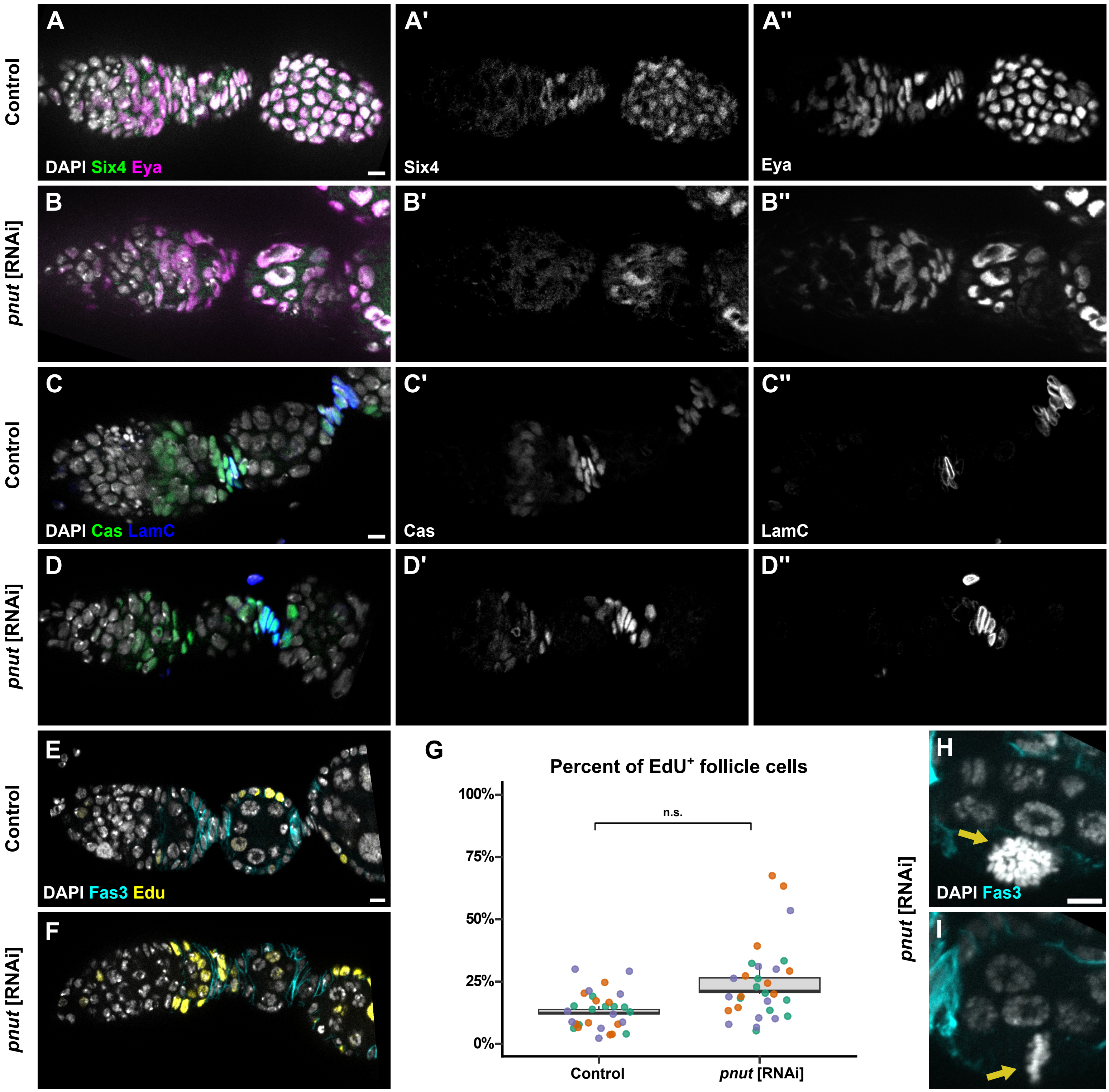

#### Supplemental Figure 5

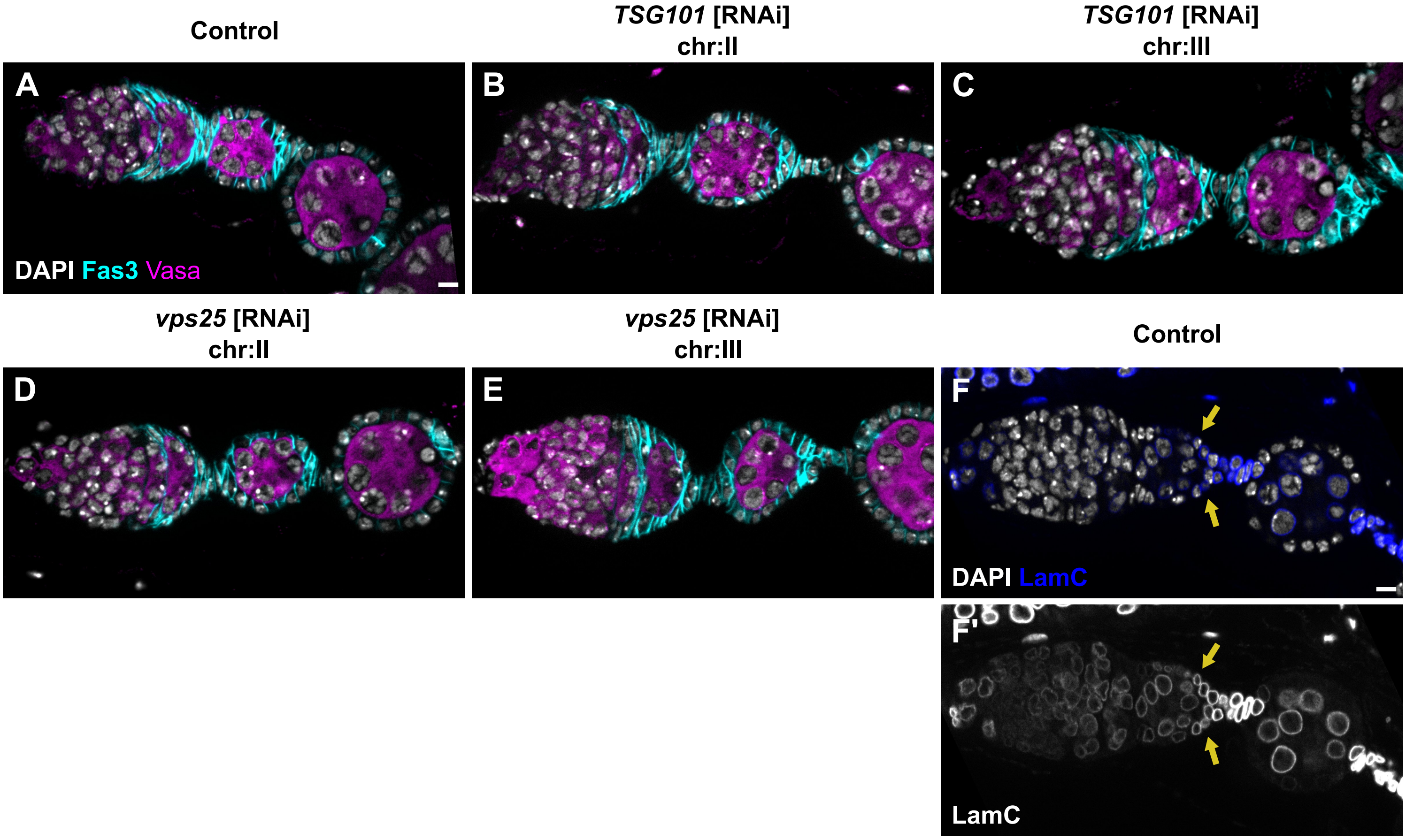
